# Perivascular space fluid contributes to diffusion tensor imaging changes in white matter

**DOI:** 10.1101/395012

**Authors:** Farshid Sepehrband, Ryan P Cabeen, Jeiran Choupan, Giuseppe Barisano, Meng Law, Arthur W Toga, the Alzheimer’s Disease Neuroimaging Initiative

**Author notes:** **Correspondence to:** Farshid Sepehrband, PhD, Laboratory of Neuro Imaging, <u>USC Mark and Mary Stevens Neuroimaging and Informatics Institute</u>, Keck School of Medicine of USC, University of Southern California, Los Angeles, CA, USA, E. Data used in preparation of this article were obtained from the Alzheimer’s Disease Neuroimaging Initiative (ADNI) database (adni.loni.usc.edu). As such, the investigators within the ADNI contributed to the design and implementation of ADNI and/or provided data but did not participate in analysis or writing of this report. A complete listing of ADNI investigators can be found at: http://adni.loni.usc.edu/wp-content/uploads/how_to_apply/ADNI_Acknowledgement_List.pdf.

## Abstract

Diffusion tensor imaging (DTI) has been extensively used to map changes in brain tissue related to neurological disorders. Among the most widespread DTI findings are increased mean diffusivity and decreased fractional anisotropy of white matter tissue in neurodegenerative diseases. Here we utilize multi-shell diffusion imaging to separate diffusion signal of the brain parenchyma from fluid within the white matter. We show that unincorporated anisotropic water in perivascular space (PVS) significantly, and systematically, biases DTI measures, casting new light on the biological validity of many previously reported findings. Despite the challenge this poses for interpreting these past findings, our results suggest that multi-shell diffusion MRI provides a new opportunity for incorporating the PVS contribution, ultimately strengthening the clinical and scientific value of diffusion MRI.

**Highlights:**

- Perivascular space (PVS) fluid significantly contributes to diffusion tensor imaging metrics
- Increased PVS fluid results in increased mean diffusivity and decreased fractional anisotropy
- PVS contribution to diffusion signal is overlooked and demands further investigation

## Introduction

Diffusion MRI is sensitive to water displacement, a physical process that is useful for characterizing structural and orientational features of brain tissue (Bihan and Breton, 1985; Le Bihan and Johansen-Berg, 2011). Diffusion tensor imaging (DTI) (Basser et al., 1994) is the most popular diffusion MRI modeling technique which has been widely used to study brain in health and disease (Alexander et al., 2007; Assaf and Pasternak, 2008; Hassan et al., 2014; Horsfield and Jones, 2002; Le Bihan et al., 2001; Sundgren et al., 2004). Over the past 30 years, many studies reported DTI-derived measures, such as fractional anisotropy (FA) and mean diffusivity (MD), in neurological diseases. A reproduced and well-known example is the observation of increased MD and decreased FA in neurodegenerative disease such as Alzheimer’s disease (Acosta-Cabronero and Nestor, 2014; Agosta et al., 2011; Amlien and Fjell, 2014; Cavedo et al., 2017; Charlton et al., 2006; Choi et al., 2005; Fellgiebel et al., 2004; Kantarci et al., 2017b, 2017a, 2014; Mayo et al., 2017; Naggara et al., 2006; Nir et al., 2013; Sexton et al., 2011; Westlye et al., 2010; Wolf et al., 2015; Zhang et al., 2009, 2007). These findings were often interpreted as the outcome of the white matter degeneration which leads to extra free space for water to displace in every direction, and therefore higher MD and lower FA (i.e. DTI findings are often interpreted as the pathological microstructural alterations of the white matter tissue). However, there remain major concerns regarding the validity of the interpretations. Neuronal degeneration is not the only occurring process, and other pathological changes related to increased glia, presence of Tau tangles and amyloid plaques (Laurent et al., 2018), may even hinder water displacement, and plausibly lower water diffusivity.

Perivascular space (PVS), also known as Virchow-Robin space, is a pial-lined, fluid-filled structure that accompany vessels entering (penetrating arteries) or leaving (draining veins) cerebral cortex (Krueger and Bechmann, 2010; Zhang et al., 1990). Due to the extensive vascularity of the brain, PVS occupies a large portion of the cerebral tissue (Osborn, 2006) (**Figure 1.a** and **1.b**). PVS volume varies across people, enlarges with aging as brain tissue shrinks, and changes in many neurological diseases (Bacyinski et al., 2017; Banerjee et al., 2017; Brown et al., 2018; Cavallari et al., 2018; Feldman et al., 2018; Kalaria, 2018; Krueger and Bechmann, 2010; Laveskog et al., 2018; Park et al., 2017). Structurally speaking, PVS has a microscopic tubular geometry that occupies extra-vascular space, with decreasing diameter as it penetrates deeper into the brain tissue. Therefore, unlike brain tissue, water molecules of PVS freely move in the microscopic scale, yet they are hindered by the vessel and the cerebral tissue in macroscopic scale. High-resolution MRI and post-mortem studies have shown that PVS in white matter has microscopic scale tubular structure with small diameter that was observed throughout the brain (Akashi et al., 2017; Bouvy et al., 2014). This morphological characteristic will result in water displacement within the white matter that can be anisotropic.

**Figure 1.**
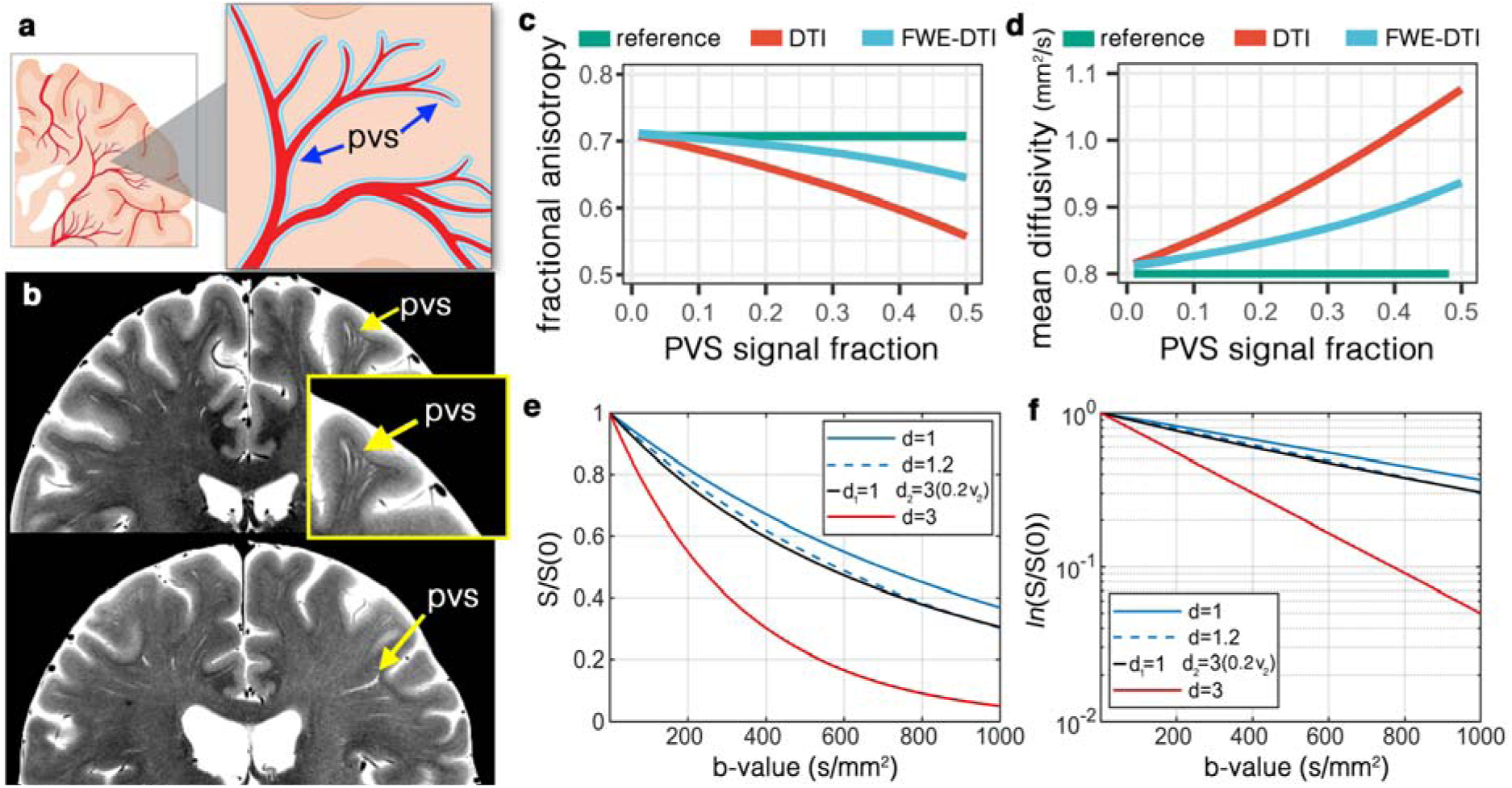
Perivascular space fluid (PVS) and its bias on diffusion tensor imaging (DTI). **(a)** Schematic view of the PVS. **(b)** High-resolution turbo spin echo images of two healthy volunteers (above: 32yr old female, below: 56yr old female), scanned at 7T. In-plane resolution of 0.3mm (interpolated to 0.15mm) was used to acquire the data (with the slice thickness of 2mm). Four averages were acquired to ensure high signal-to-noise ratio. Note that PVS presents throughout the white matter, with larger diameter PVSs closer to the cortex and smaller diameter as it penetrates deep into the white matter. Systematic bias from PVS on DTI was simulated for the fractional anisotropy **(c)** and mean diffusivity **(d)**. As PVS increases the amount of bias amplifies. Free-water elimination (DTI-FWE) technique is also included, which is also affected by PVS presence but to a smaller extent. A plot of diffusion MRI signal for 3 different examples of mean diffusivity (d=1 μm^2^/ms, blue line; d=1.2 μm^2^/ms, blue dashed line; and d=3 μm^2^/ms, red line) is illustrated in **(e)** and the log of the signal is plotted in **(f)**. Note that an increased mean diffusivity from 1 μm^2^/ms to 1.2 μm^2^/ms has a similar signal profile as a scenario with no increased diffusivity but 20% of PVS presence (i.e. PVS signal fraction of 0.2; black line).

When DTI measures are estimated, the derived measure reflects the diffusion properties of both the tissue and fluid from the PVS. Given the relatively fast diffusivity of the water in PVS, even a small portion of PVS in an imaging voxel can have a substantial effect on the voxel averaged DTI measures, due to the partial volume effect (Alexander et al., 2001). Here we focus on the effect of PVS fluid on DTI-derived measures, namely FA and MD. Our experiments demonstrate that a failure to incorporate this fluid compartment can impose a systematic bias in how DTI can be interpreted. We show that in a brain tissue with a given DTI characteristic, if the amount of PVS increases (**Figure 1.c**), DTI modeling would result in an increased MD and decreased FA. While disrupted tissue microstructure is often cited when changes in DTI parameters are observed, the bias imposed by changes in PVS fluid provides a hypothetical but compelling alternative explanation for many reported findings in DTI studies of neurodegenerative diseases.

## Method

In order to assess this bias we considered two tensors in a voxel (Pierpaoli and Jones, 2004), one for the tissue and the other for the PVS compartment. This model was utilized because it separates PVS and the tissue and allows the examination of the effect of PVS fluid on DTI signal. It also enables direct comparison between tensor-derived measures across compartments. For clarity, the images relating to the tissue are called tissue tensor images (TTI), which were used to investigate the bias of the DTI. DTI reflects the *voxel values,* but TTI reflects *tissue values* and are referred to accordingly. Experimental data showed that an anisotropic model of the PVS fits better to diffusion data compared to an isotropic model (described below). Therefore, throughout this study we used an anisotropic model of PVS as the reference model, when evaluating DTI measures. It should be noted that a diffusion MRI acquisition with multiple b-values is required (multi-shell diffusion MRI) (Pierpaoli and Jones, 2004) for multi-compartment modeling of brain tissue, and datasets were selected accordingly.

### Experimental data to asses PVS anisotropy

To ensure that PVS diffusion signal is anisotropic, we acquired a multi-shell non-conventional diffusion MRI of a healthy 32-years-old female volunteer and assessed the goodness of fit of an anisotropic model versus an isotropic model. An hour of scan was conducted to acquire 632 diffusion MRI volumes. Multi-shell diffusion MRI with b-values of 0, 200, 400, 600, 800, 1000, 1200, 1500 and 2000 s/mm^2^ was acquired with isotropic resolution of 1.5 mm^3^ using a 3T scanner (Prisma, Siemens Healthcare, Erlangen, Germany), with an acquisition sequence similar to Human Connectome Project (HCP) (Essen et al., 2013). Thirty gradient-encoding directions for low b-value shells (<1500) and 60 gradient-encoding directions for other shells were acquired in both anterior-posterior and posterior-anterior phase encoding directions. We used a single-channel quadrature transmit radiofrequency (RF) coil and a 32-channel receive array coil (Nova Medical Inc., MA). In addition to diffusion MRI, high-resolution T2-weighted images were also acquired to locate PVS in fine detail to aid spotting regions with high PVS presence. T2-weighted images using turbo-spin echo sequences with in-plane resolution of 340 μm (interpolated to 170 μm) and 2 mm slice thickness were collected with two averages and two concatenations. The institutional review board of the University of Southern California approved the study. Informed consent was obtained from the volunteer, and the image datasets were anonymized.

*dcm2nii* was used to convert the *dicom* images to the *nifti* file format (Li et al., 2016). Diffusion MRI data were corrected for subject motion, eddy current, EPI distortion. FSL’s TOPUP was used to correct for B0-inhomogeneity distortion using two opposing phase encoded images (Andersson et al., 2003). FSL’s EDDY was used to correct for current induced field inhomogeneity and subject’s head motion (Andersson et al., 2012), followed by correction for the gradient nonlinearity. Two bi-tensor models were fitted to the diffusion data, one allowing anisotropic diffusion for PVS and one constrained to isotropic diffusion. Quantitative Imaging Toolkit (QIT) (Cabeen et al., 2018) was used for fitting. Except for the diffusion profile of the PVS compartment, an identical fitting routine was used for both models. Fitting was performed using constrained trust-region derivative-free optimization using Powell’s BOBYQA algorithm (Powell, 2009). The signal fraction was required to be between zero and one, the fluid compartment was required to be axially symmetric fluid compartment with positive diffusivities and have an axis aligned to the tissue principal direction, and the tissue compartment was constrained to be positive definite using a re-parameterization with the Cholesky decomposition. Models were compared by comparing the root mean square error of the fit and also by performing the Akaike information criteria (AIC) test (Akaike, 1974), as described here (Burnham and Anderson, 2004).

As expected, the diffusivity of the PVS compartment was not isotropic in white matter (**Figure 2**) and therefore diffusion profile of the PVS compartment was not fixed to an isotropic profile. Anisotropic model fitted more accurately to the white matter voxels compared to the isotropic model (**Figure 2.e** and **2.f**). The fitting was particularly superior in voxels with high PVS presence. For example, pre-cortical white matter voxels around centrum semiovale were best modeled when the anisotropic model of the PVS was utilized. Statistically, the root mean square error of the anisotropic fit was significantly lower than that of the isotropic model (*t*(15095) = 147.73, *p* < 0.0001). AIC test resulted to the same conclusion, in which the anisotropic model of the PVS compartment outperformed the isotropic model (AIC score was significantly lower in the anisotropic model: t(15095) = −142.15, *p* < 0.0001). An isotropic assumption for the PVS compartment is evidently not optimal. Therefore, throughout this study we used an anisotropic model of PVS, as the reference model to evaluate against DTI measures.

**Figure 2.**
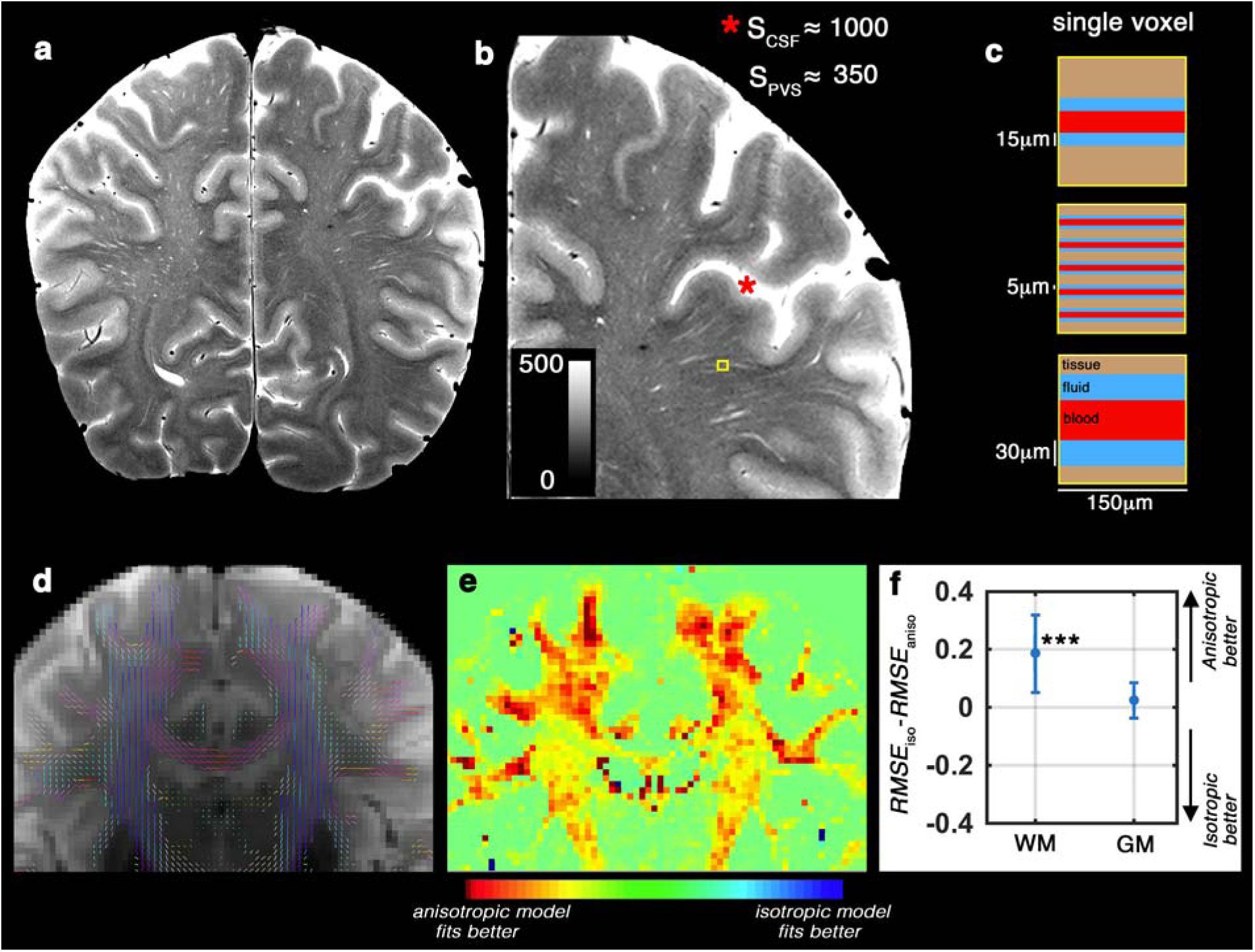
Diffusion of the perivascular space fluid is anisotropic in conventional DWI. **(a)** T2-weighted image of a healthy 32 years old volunteer, scanned at 7T with in-plane resolution of 150 μm^2^. **(b)** Zooming into a region with high perivascular space (PVS) presence. Signal value of the cerebrospinal fluid (CSF) voxels were much higher than the hyperintense PVS voxels, because PVS voxels partially share the imaging signal with white matter and vessel (Color range is fixed to [0 500], to better visualize the PVS and avoid CSF saturation). Schematic representation of the PVS partial volume for three different possible scenarios are presented in **(c)**. T2-weighted image (unweighted diffusion image of the same subject, scanned at 3T) is shown in **(d)** and the DTI-derived tensor glyphs of the white matter voxels are overlaid on top. **(e)** shows the spatial distribution of the fitting error difference (Akaike information criteria scores resulted to a similar heat map) between anisotropic and isotropic models of the PVS fluid diffusion. Note that the highest differences are observed in white matter voxels near cortex, with high PVS presence. The quantitative difference is presented in **(f)**. Same results as (**e** and **f**) was obtained from Akaike information criteria test. Anisotropic model fitted better to the data, particularly within the white matter.

### Simulation

In the simulation experiments, the diffusion-weighted signal was synthesized with biologically plausible tissue and perivascular space (PVS) contributions. The synthetic model had a baseline signal of one, a fluid signal fraction of 0.1, and axially symmetric and aligned tensors for the tissue and fluid compartments. The tissue compartment had an axial diffusivity of 1.6 μm^2^/ms and a radial diffusivity of 0.4 μm^2^/s. The fluid compartment had an axial diffusivity of 3 μm^2^/s and a radial diffusivity of 2.5 μm^2^/s. The signal was simulated using a multi-shell acquisition scheme. Diffusion encoding gradients were optimized for multi-shell sampling, using the *q-space sampling Web application* (http://www.emmanuel-caruyer.com/q-space-sampling.php) (Caruyer et al., 2013). Total of nine shells, with b-values ranging from 0 s/mm^2^ to 2000 s/mm^2^ were used (at 250 s/mm^2^ steps, with 90 q-space sampling per shell). Noise with a standard deviation of 0.025 was added to the simulated signal, then diffusion tensor imaging (DTI) (Basser et al., 1994; Bihan and Breton, 1985) and DTI free water elimination (DTI-FWE) (Pasternak et al., 2009) models were fitted to the data, and finally diffusion parameters were extracted from the fitted models.

### High-resolution 7T images

High-resolution T2-weighted images were acquired to visualize PVS in fine detail. Two healthy adult females (32 and 56 years old) were scanned on a 7 Tesla (7T), whole-body scanner (Terra, Siemens Healthcare, Erlangen, Germany) using a single-channel quadrature transmit radiofrequency (RF) coil and a 32-channel receive array coil (Nova Medical Inc., MA). The institutional review board of the University of Southern California approved the study.

Informed consent was obtained from the volunteers, and the image datasets were anonymized.

T2-weighted using turbo-spin echo sequences with in-plane resolution of 300 μm (interpolated to 150 μm) and 2 mm slice thickness were collected. Four averages and two concatenations were acquired to enhance image SNR and CNR (Sepehrband et al., 2018). With echo time of 73 ms, repetition time of 3.5 s and total of 25 slices, the acquisition time was 12 minutes.

### HCP data

We evaluated the effect of PVS on DTI measures on a large cohort of young healthy adults, in whom pathological white matter fluid such as microcysts and lacunar infarcts are not expected. We also focused on voxels and regions were PVS presence could be confirmed from structural MRI. We downloaded structural and diffusion magnetic resonance imaging (MRI) data provided by the HCP (Essen et al., 2013), namely “S900 release”. This dataset includes 861 healthy participants (age, 22–35 years) with multi-shell diffusion MRI (1.25 mm^3^ resolution) and structural T1-weighted and T2-weighted images (0.7 mm^3^ resolution images), suitable for our analyses. The diffusion MRI image included three shells of b-values (1000, 2000 and 3000 s/mm^2^), each with 90 diffusion-weighted images. In addition, 18 non-diffusion-weighted images were acquired. FA and MD were compared in this cohort with and without considering the PVS contribution.

### HCP data analysis

We used preprocessed data using methods detailed previously, which were preprocessed using HCP pipelines (Glasser et al., 2013; Milchenko and Marcus, 2013; Sotiropoulos et al., 2013). In brief: the structural images were corrected for gradient nonlinearity, readout, and bias field; aligned to AC-PC “native” space and averaged when multiple runs were available; then registered to MNI 152 space using FSL (Jenkinson et al., 2012)’s FNIRT. The native space images were used to generate individual white and pial surfaces (Glasser et al., 2013) using the FreeSurfer software (Fischl, 2012) and the HCP pipelines (Glasser et al., 2013; Sotiropoulos et al., 2013). FSL’s TOPUP was used to correct for B0-inhomogeneity distortion using two opposing phase encoded images (Andersson et al., 2003). FSL’s EDDY was used to correct for current induced field inhomogeneity and subject head motion (Andersson et al., 2012), followed by correction for the gradient nonlinearity. Diffusion data were registered to the structural T1-weighted AC-PC space using the non-diffusion-weighted volume. The diffusion gradient vectors were rotated accordingly.

DTI, DTI-FWE and tissue tensor imaging (TTI) models were fitted to HCP subjects using Quantitative Imaging Toolkit (QIT) (Cabeen et al., 2018). For a robust estimation of DTI measure, the shell with the b-value of 1000 s/mm^2^ was separated and the tensor model was fitted to each voxel of the volume using a non-linear least square fitting routine. DTI-FWE and TTI were fitted to the complete diffusion data. DTI-FWE model fitting was performed using a custom implementation of the procedure described by Hoy *et al.* (Hoy et al., 2014), in which the fluid compartment is assigned a constant diffusivity of 3 μm^2^/s and the optimal signal fraction parameter is determined through a grid search with linear least squares of the tissue tensor compartment at each grid point. The TTI model fitting was performed similar to the fitting of the experimental data. TTI fitting was initialized with the parameters obtained from the DTI-FWE model, and TTI parameters were constrained as follows: the signal fraction was required to be between zero and one, the fluid compartment was required to be axially symmetric fluid compartment with positive diffusivities and have an axis aligned to the tissue principal direction, and the tissue compartment was constrained to be positive definite using a re-parameterization with the Cholesky decomposition.

Diffusion MRI-derived measures were compared in different areas of the white matter: in voxels with high PVS signal fraction and then in four atlas-driven regions of the white matter that are known to have varying PVS appearance in healthy adults (Osborn, 2006), namely: corpus callosum (low PVS appearance), para-hippocampus (intermediate PVS appearance), centrum semiovale (high PVS appearance), and superior-frontal part of the white matter (an additional randomly selected region). White matter voxels with high PVS appearance were selected from high-resolution structural images. We noted that the “T1-weighted divided by T2-weighted” images, provided as part of the HCP release, can clearly highlight voxels with high PVS presence. The additional clarity is because fluid appears hyperintense in T2-weighted images and hypointense in T1-weighted images. A threshold of 2.5 (based on manual inspection of the voxels with high PVS presence) was used to segment PVS, where a voxel with “T1-weighted divided by T2-weighted” intensity of smaller than 2.5 was considered a voxel with high PVS presence (**Figure 3.a** is a given example). An inflated mask of lateral ventricles was then used to excluded incorrectly segmented voxels in the periventricular areas, mainly observed in the body and posterior horn of the ventricles. Four white matter regions were extracted from FreeSurfer’s white matter segmentation outputs (Fischl, 2012), which were derived using Desikan-Killiany atlas (Desikan et al., 2006). When comparing diffusion MRI-derived measures, paired t-test and Pearson correlation were used.

**Figure 3.**
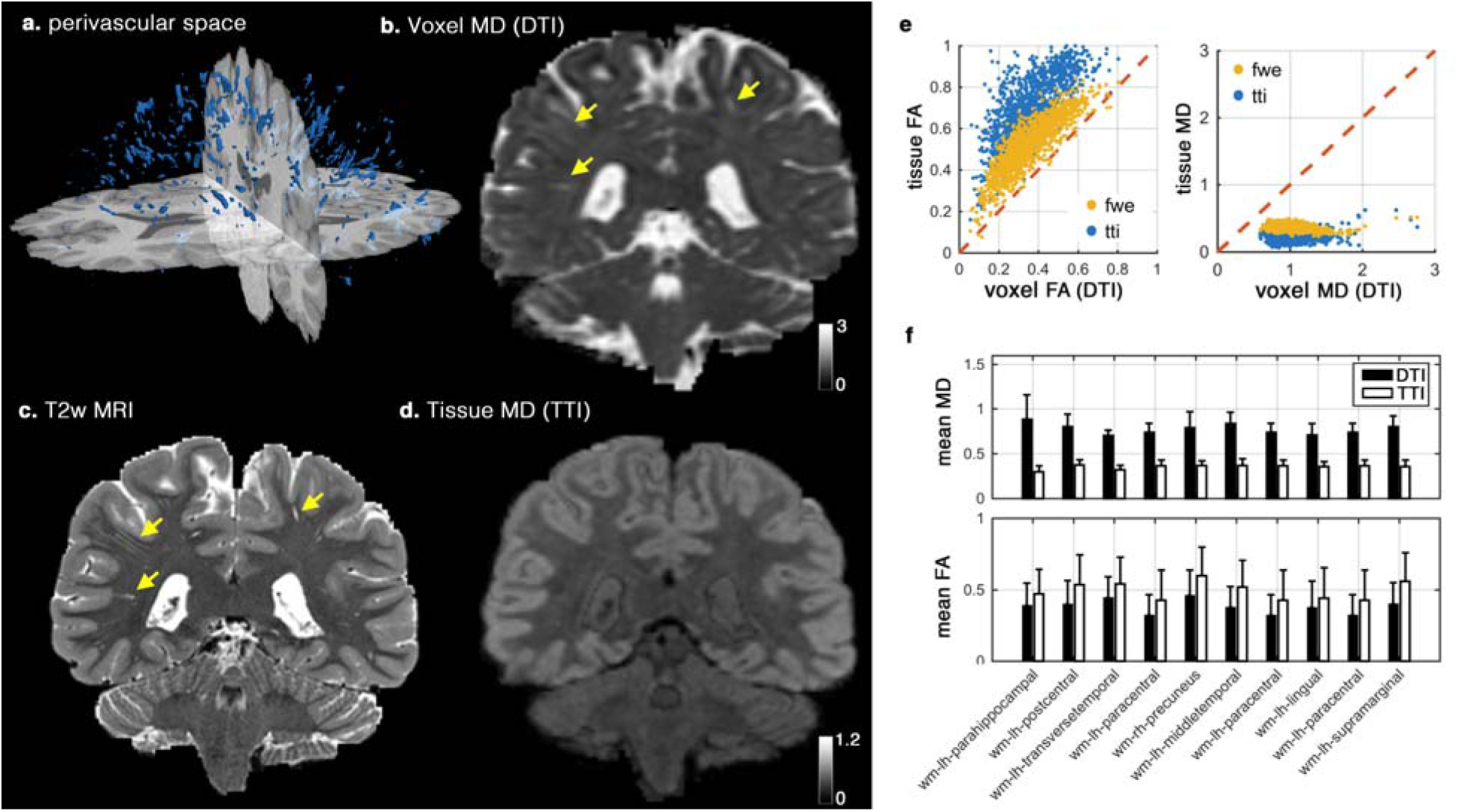
Investigating the effect of the perivascular space (PVS) on DTI in single subject level. Segmented PVS voxels of a healthy subject from human connectome project is plotted in **(a)**. Voxel mean diffusivity (MD) derived using DTI, T2-weighted image, and Tissue MD derived from tissue tensor imaging (TTI) are plotted **(b-d)**, respectively. Note that the expected white matter homogeneity is preserved in tissue MD map, while voxel MD is largely affected by the PVS contribution (extreme cases are demonstrated by yellow arrows). Correlation of the voxel values and the tissue values that were derived from DTI and TTI are plotted in **(e)**. Values from free water elimination (FWE) technique are included for comparison. Mean values of MD and FA from DTI and TTI across 10 white matter regions are also plotted **(f)**. All differences of MD and FA mean values in **(f)** are significant at *p*<0.001 (using a paired *t*-test). Diffusivity values arein μm^2^/ms.

### ADNI-3 data

Data used in the preparation of this article were obtained from the Alzheimer’s Disease Neuroimaging Initiative (ADNI) database (http://adni.loni.usc.edu). The ADNI was launched in 2003 as a public-private partnership, led by Principal Investigator Michael W. Weiner, MD. The primary goal of ADNI has been to test whether serial MRI, positron emission tomography (PET), other biological markers, and clinical and neuropsychological assessment can be combined to measure the progression of mild cognitive impairment (MCI) and early Alzheimer’s disease (AD).

The PVS bias was investigated on an Alzheimer disease neuroimaging initiative 3 (ADNI-3) cohort (Weiner et al., 2017), in which multi-shell diffusion MRI data is available. Data of 62 subjects with multi-shell diffusion MRI was downloaded from the ADNI database (http://adni.loni.usc.edu) (Toga and Crawford, 2010). One young CN subject (54-year-old) was excluded. Subjects were divided into two groups of cognitively normal (CN) subjects (*N*=37, 24 females) and MCI patients (*N*=24, 7 females). Average age of the CN (*M*=73.7, *SD*=7.9) and the MCI group (*M*=75.5, *SD*=6.8) were not statistically different (t(59)=0.93, *p*=0.36). The MCI group consisted of patients with the following cognitive stages: significant memory concerns (*N*=2), early MCI (*N*=7), MCI (*N*=12), late MCI (*N*=3). FA and MD were compared between the CN and MCI groups, with and without considering the fluid contribution.

### ADNI-3 data analysis

All ADNI-3 images used in this study were acquired using Siemens Prisma or Prisma_fit 3T scanner (Siemens Healthcare, Erlangen, Germany), on six different sites, using a standardized diffusion MRI sequence (Wyman et al., 2013). Diffusion MRI data was acquired using the following parameters: 2D echo-planar axial imaging, with sliced thickness of 2mm, in-plane resolution of 2mm^2^ (matrix size of 1044 x 1044), flip angle of 90°, 126 diffusion-encoding images with three b-values (6 directions for b-value=500 s/mm^2^, 48 directions for b-value=1000 s/mm^2^, 60 directions for b-value=2000 s/mm^2^), with 13 non-diffusion-weighted images were acquired.

After downloading the raw images, *dcm2nii* was used to convert the *dicom* images to the *nifti* file format (Li et al., 2016). Diffusion MRI were corrected for eddy current distortion and for involuntary movement, using FSL TOPUP and EDDY tools (Andersson et al., 2012, 2003). DTI, DTI-FWE, and TTI models were fitted using the same procedure as with the HCP data. Data was analyzed using QIT to examine diffusion tensor parameters in deep white matter, as defined by the Johns Hopkins University (JHU) white matter atlas (Mori et al., 2008). The JHU regions were segmented in each scan using an automated atlas-based approach described in Cabeen et al., (Cabeen et al., 2017) in which deformable tensor-based registration using DTI toolkit (DTI-TK) (Zhang et al., 2006) was used to align the subject data to the Illinois institute of technology (IIT) diffusion tensor template (Zhang et al., 2011), and subsequently to transform the JHU atlas regions to the subject data and compute the average of each diffusion tensor parameter with each JHU region.

We used linear regression when investigating the relation between diffusion-derived measures with the cognitive stage using an ordinary least square fitting routine, implemented with the statsmodels.OLS module in Python 3.5.3 (StatsModels version 0.8.0 – other Python packages that were used are Pandas version 0.20.3 and NumPy version 1.13.1). Multiple regressions were fitted to regional mean values, one region at a time. For every instance, sex, estimated total intracranial volume, and age were included as covariates. The Benjamini–Hochberg procedure with a false discovery rate of 0.1 was used to correct for multiple comparisons. Diffusion MRI-derived measures were compared using paired t-test and Pearson correlation. Bland-Altman plots (Altman and Bland, 1983) were used to investigate whether DTI and TTI were systematically different. Bland-Altman plot analysis are designed to investigate a bias between the mean differences (Bland and Altman, 1995), where a distribution above or below 0 (on the y-axis) indicates a bias.

## Results

### Simulation data

Diffusion MR signal in a white matter voxel was simulated by changing the amount of PVS. Simulation experiments show how FA and MD of the tissue, when modeled using DTI, deviate from the tissue ground truth values, as the amount of PVS increases (**Figure 1.e** and **1.f**). For example, a 20% increase in PVS signal contribution would result in a same signal change if the tissue MD increases from 1 to 1.2 μm^2^/ms. Our simulations show that even a model incorporating an isotropic free water compartment, i.e. without incorporating fluid anisotropy, could still systematically bias results in the same direction as the DTI bias. However, the scale of this bias is significantly lower (**Figure 1.c** and **1.d**).

### DTI bias in healthy subjects

We demonstrated the influence of the PVS on the DTI-derived maps first on a single subject and then investigated it on a large cohort of 861 healthy subjects. Subject-level investigation showed that the MD map of DTI was significantly affected by PVS contribution (**Figure 3.a-d**). Incorporating PVS contribution has dramatically improved the clarity of the MD map (**Figure 3.d**), wherein white matter homogeneity is preserved. Also, the white-gray matter contrast is greater compared with DTI-derived MD. The PVS map visually resembles the T2-weighted image, without the PVS contrast (more examples are provided in **Supplemental Figure 1** and **2**). We observed that ignoring PVS contribution to diffusion MRI signal could even influence the visual presentation of the FA map, particularly around PVS area (**Supplemental Figure 3**).

Quantitative MD and FA values from DTI were significantly different from TTI, showing the expected systematic bias of increased MD and decreased FA. **Figure 3.e** shows the correlation of TTI-derived MD and FA with DTI-derived MD and FA. MD values from DTI were significantly higher (*t*(1345) = 114.70, *p* < 0.001) than tissue MD from TTI. Tissue MD values were more stable compared with voxel MD values 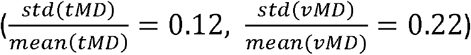, reflecting the expected white matter quantitative homogeneity. FA values from DTI were significantly lower in PVS voxels (*t*(1345) = —119.55, *p* < 0.001). Tissue FA and voxel FA were highly correlated (*r* = 0.87, *n* = 1345, *p* < 0.001), but MD values were weakly correlated (*r* = 0.15, *n* = 1345, *p* < 0.001). The quantitative difference was observed beyond the PVS voxels and showed to affect the regional values (**Figure 3.f**). Values of all WM regions were significantly different between DTI and TTI (all differences were significant at *p* < 0.001 level).

To determine whether this effect is generalized across individuals, we characterized the DTI bias in a typical adult population by quantitatively examining 861 healthy subjects from human connectome project (HCP) (**Figure 4**) (Essen et al., 2013). Empirical results confirmed the simulation study, where an increased MD and a decreased FA in DTI results were observed. Values were compared in PVS voxels and in white matter regions with different expected concentration of PVS, namely: corpus callosum, para-hippocampus, centrum semiovale, and superior-frontal part of the white matter (**Figure 4.c** and **4.f**). When PVS voxels were looked at, a large and significant difference between TTI and DTI measures were observed (*t*(860) = 283.13, *p* < 0.001). Voxel MD values from DTI (*mean* = 0.94, *std* = 0.07) were almost three times higher than TTI values (*mean* = 0.39, *std* = 0.02). This is not surprising given that much of the volume of these voxels are occupied by PVS. MD and FA were also significantly different (all at *p* < 0.001 level) even at region-averaged level across DTI, DTI-FWE and TTI (**Figure 4.c** and **4.f**). FA values derived from DTI-FWE were closer to DTI, but MD values derived from DTI-FWE were closer to TTI, both significantly different (both at *p* < 0.001 level), confirming that DTI-FWE is not a remedy to the PVS imposed bias (see detailed statistics in **Supplemental Note 1**).

**Figure 4.**
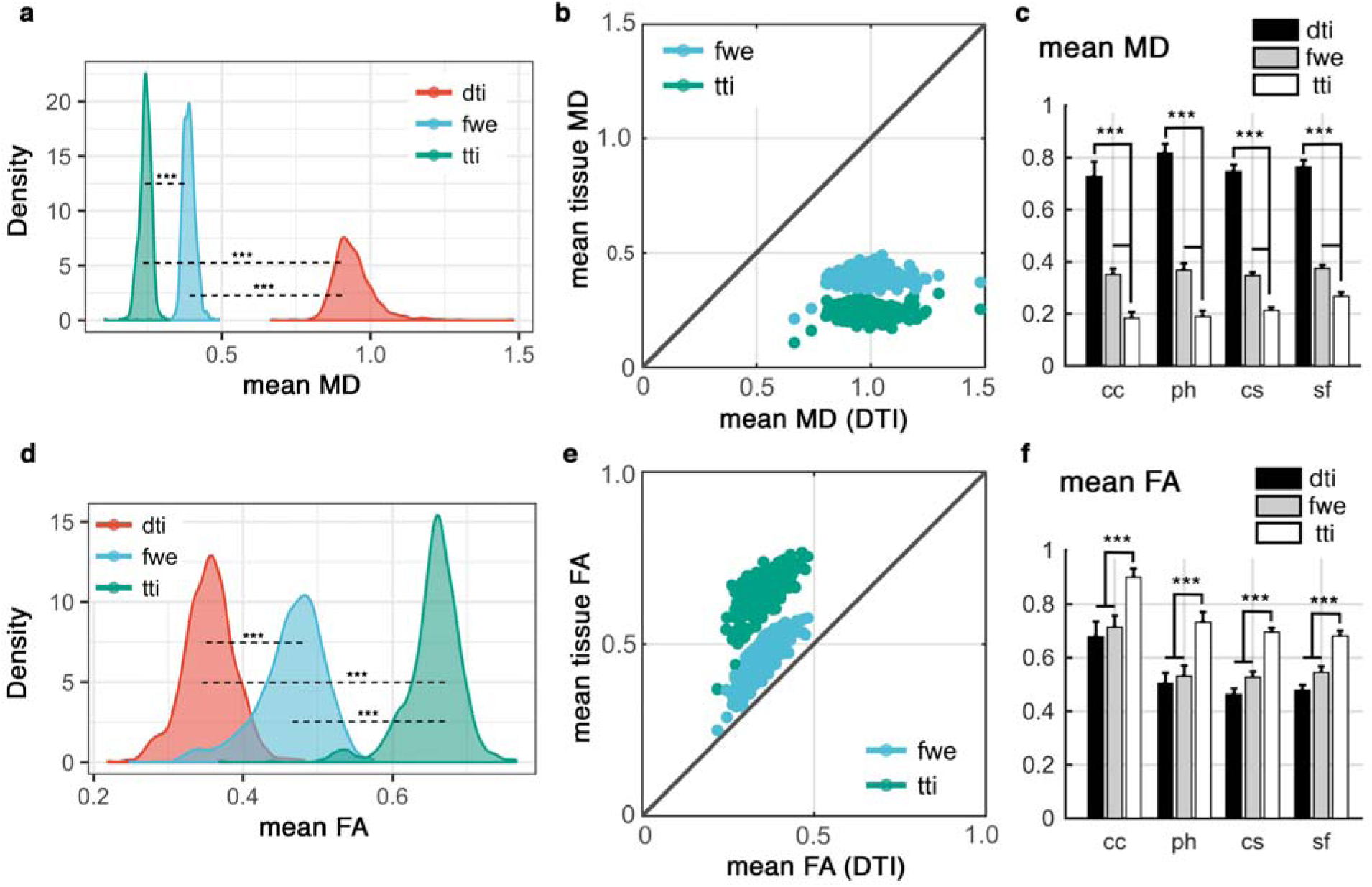
Quantitative investigation of the influence of perivascular space (PVS) on DTI, across 861 subjects of the human connectome project data. Values from DTI were compared with those from free water elimination (FWE) and tissue tensor imaging (TTI) techniques. Mean MD of the PVS voxels are compared in **(a)** and **(b)**. Mean MD values were compared across four white matter regions from Desikan-Killiany atlas **(c)**. Regions are: corpus callosum (cc), para-hippocampus (ph), centrum semiovale (cs) and superior-frontal part of the white matter (sf). All differences are corroborating simulation and subject-level results and are significant at *p*<0.001 (using paired t-test). Similar investigation on the fractional anisotropy (FA) values are shown in **(d-f)**. Diffusivity values are in μm^2^/ms.

### DTI bias in a neurodegeneration study

We examined how the PVS DTI bias affects the study of neurodegeneration using data from the ADNI-3 project (Weiner et al., 2017). MD and FA values from DTI were significantly different than those from TTI across all regions of the white matter (20 random regions are plotted in **Figure 5.e** and **5.f**, and the complete list is presented in **Supplemental Figure 4**). The differences were similar to the expected bias and in corroboration with simulation and HCP data analysis. The magnitude of the bias was larger across MD measures compare to FA. The Bland-Altman plots confirm that measures from DTI and TTI are systematically different (MD of DTI is higher, and FA of DTI is lower).

**Figure 5.**
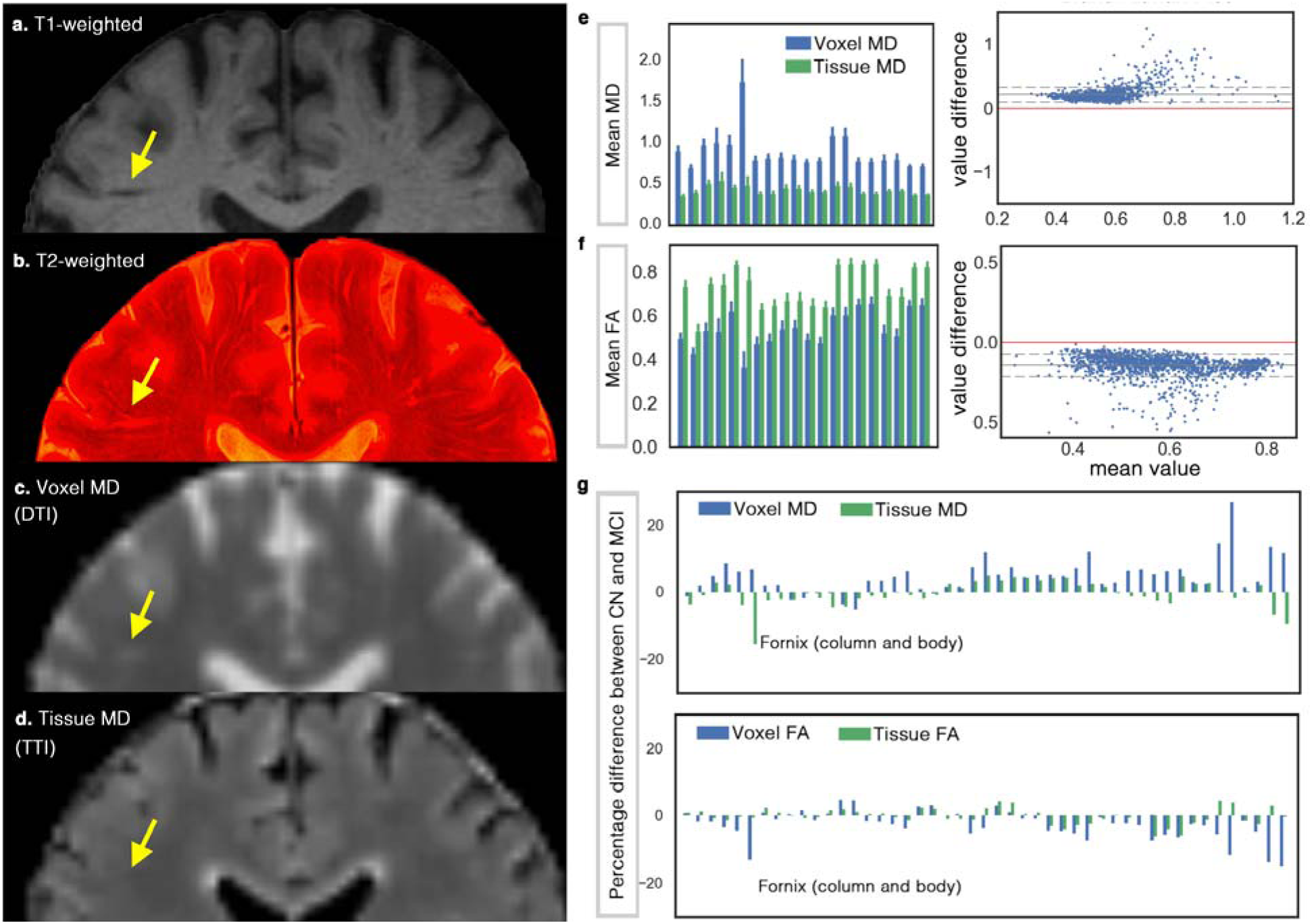
Investigating the effect of perivascular space (PVS) on DTI measures on a cohort of ADNI-3, including 37 cognitively normal (CN) subjects and 24 mild cognitively impaired (MCI) patients. **(a)** shows the T1-weighted, in which a portion of the PVS can be seen. By superimposing high-resolution T2-weighted image on the T1-weighted image **(b)** more PVS are detectable. Mean diffusivity (MD) map from DTI and tissue tensor imaging (TTI) are mapped in **(c)** and **(d)**, respectively. Note that MD from TTI preserved the expected white matter homogeneity by separating the PVS water from the tissue, yet successfully mapped the white matter hyperintensities. White matter hyperintensities are indistinguishable in the MD map from DTI, given the partial volume effect of the periventricular space. Mean voxel MD and tissue MD of 20 randomly selected (for the sake of space) regions from John Hopkins white matter atlas are shown in **(e)**. Bland-Altman plot was drawn to compare DTI versus TTI, which shows the systematic bias of the DTI measures (note that if two techniques were equal the values would show a standard deviation around the value difference of zero, shown by red), **(f)** shows a similar analysis for the fractional anisotropy (FA) measures. The complete chart of John Hopkins white matter regions is illustrated in **Supplemental Figure 4. (g)** compares DTI results with TTI when CN group was compared with MCI. Note that for MD, most regions show increased MD (as reported in the literature), but when TTI was used, the inverse pattern was observed in many cases. Fornix is an extreme case of this example (while a large difference in Fornix was observe, it was not statically significant). Diffusivity values are in μm^2^/ms.

An example group-level analysis was performed on the ADNI-3 data to judge if the findings differ when we incorporate a PVS contribution. When comparing MD values of CN and MCI groups using DTI, twenty-two regions were significantly different after correcting for multiple comparison using Benjamini–Hochberg procedure (Kwee and Kwee, 2007) and with the false discovery rate of 0.1 (age, sex and brain size were included in the regression). These regions and the statistics are reported in **Supplemental Note 2.** When TTI was used, or when the signal fraction of the PVS was included in the regression, none of those regions were significant.

When comparing FA values of CN and MCI groups using DTI, twelve regions were significantly different after correcting for multiple comparison using Benjamini-Hochberg procedure and with the false discovery rate of 0.1 (age, sex and brain size are included in the regression). Most of these regions shown no significant association when TTI was used or when PVS signal fraction was included in the DTI regression model, except for two regions that were significantly different: **1.** Hippocampal connection of the right cingulum (*p* < 0.001, *FDR*(*q*) = 0.0021), **2.** The Stria terminalis of the right fornix (*p* = 0.0032, FDR(*q*) = 0.0042). The former region was not significantly different between studied groups when a more conservative false discovery rate was applied (i.e. false discovery rate of 0.05).

## Discussion

Here we showed that ignoring PVS fluid can systematically bias DTI findings. This bias affects how DTI-derived measures such as MD and FA are interpreted. An increased MD or decreased FA could be due to a physiologically normal higher amount of PVS fluid in the voxel. It could also be pathological (for example, PVS enlargement). Hence, ignoring this compartment negatively affects the mechanistic power of diffusion MRI. We also showed that employing a multi-shell acquisition strategy enables compartmentation of the diffusion signal to PVS and parenchyma, providing additional insight into diffusion signal change. Such capability makes diffusion MRI a powerful tool to assess the mechanistic changes underlying white matter physiological and pathological changes.

In order to assess the effect of PVS on DTI measures, we used a bi-tensor model to separate PVS signal from tissue signal, similar to (Pierpaoli and Jones, 2004), but by incorporating biological prior knowledge about PVS fluid diffusion profile. We considered an anisotropic water diffusivity for the PVS compartment that is aligned with white matter tracts. We also assumed that diffusivity of the PVS fluid is higher than white matter diffusivity, but not fixed. This prior knowledge about PVS fluid aids a robust fitting of the bi-tensor model to diffusion data, which is otherwise an ill-conditioned fitting problem. Both our experimental data and literature support these assumptions.

Doucette et al, recently showed that spin echo perfusion dynamic susceptibility contrast signal depends on white matter fiber orientation, which is due to vessels running in parallel with white matter tracts (Doucette et al., 2018). They showed that only a model that assumes a high diffusion coefficient (i.e. PVS) around the vessels is able to fit the data (Doucette et al., 2018; Hernández-Torres et al., 2017). In addition, histology studies exhibit the anisotropy of the vascular architecture and also showed that their caliber can widely vary (Amato et al., 2016; Cavaglia et al., 2001; Duvernoy et al., 1981). We also provide experimental evidence that the diffusivity of the PVS in white matter is anisotropic. PVS fluid is hindered by tissue parenchyma and vessel wall, and the capillary flow (Le Bihan, 1990) could selectively and non-linearly affect the diffusivity of PVS, suggesting that a fixed diffusivity may not be an optimum choice.

Several techniques and previous studies have included free water in diffusion tensor modeling (Berlot et al., 2014; Hoy et al., 2017; Metzler-Baddeley et al., 2012; Pasternak et al., 2009; Pierpaoli and Jones, 2004) or aimed to eliminate it by modifying the imaging sequence (Papadakis et al., 2002), to address the CSF partial volume effect in white matter boundaries (e.g. near ventricle). Most of these studies used a fixed-diffusivity isotropic diffusion model and/or treated PVS fluid as a factor to eliminate. Non-zero volume fraction of the fluid compartment in these techniques has been assumed to relate to extra-cellular fluid. Here in addition to the introduction and examination of this systematic of DTI measures, we also emphasize that Efforts to eliminate fluid contributions may not be the right approach, as parameters obtained from this compartment could be an imaging signal of significant scientific value. For example, Taoka *et al.* recently showed that diffusivity along the perivascular space may reflect impairment of the glymphatic system (Taoka et al., 2017). The extent to which these findings may be affected by the choice of the model is yet to be examined. More recently, Thomas *et al.* showed that DTI measures fluctuates during the day, which could be a reflection of physiological changes of the glymphatic system, including changes in the PVS fluid amount (Thomas et al., 2018).

Some previous approaches for free water elimination have used single shell data (Pasternak et al., 2009); however, our work shows the importance of more complex parameterization of the fluid compartment that requires a multi-shell diffusion acquisition, similar to (Hoy et al., 2014; Pasternak et al., 2012). It is plausible that such single shell free water elimination techniques may also be biased in the presences of anisotropic PVS fluid, but this remains an open question to be investigated further.

PVS can be mapped with high-resolution T2-weighted imaging only in some voxels where PVS contribution is above the contrast-to-noise ratio (Kwee and Kwee, 2007). It can however be fully mapped and quantified with diffusion MRI by extracting the signal fraction of the PVS (**Supplemental Figure 1** and **2**). The signal fraction of the PVS and its diffusion characteristics are valuable measures with great potential clinical significance. Microscopic or mesoscopic tissue degeneration may result in microscopic and mesoscopic tissue shrinkage which could change the MRI appearance of the surrounding PVS. Our experiments suggest that diffusion MRI opens a window to characterizing this potentially significant tissue alteration *in vivo.*

### An example between-group study (ADNI-3)

Results from ADNI-3 subjects with multi-shell diffusion MRI data were in-line with simulation and HCP data results (**Figure 5**). It should be noted that voxels of the ADNI-3 diffusion MRI data are 4.1 times larger than that from HCP data, yet the influence of the PVS on the MD map was clearly apparent (**Figure 5.c** and **5.d**). Interestingly, the tissue MD map not only separated PVS from the tissue, but also was able to map the periventricular white matter hyperintensities. MD map of the TTI resembles the FLAIR contrast, with the advantage of having additional quantitative value. To highlight the inter-group variability of the PVS concentration, one example image from CN and MCI groups is depicted in **Figure 6.**

**Figure 6.**
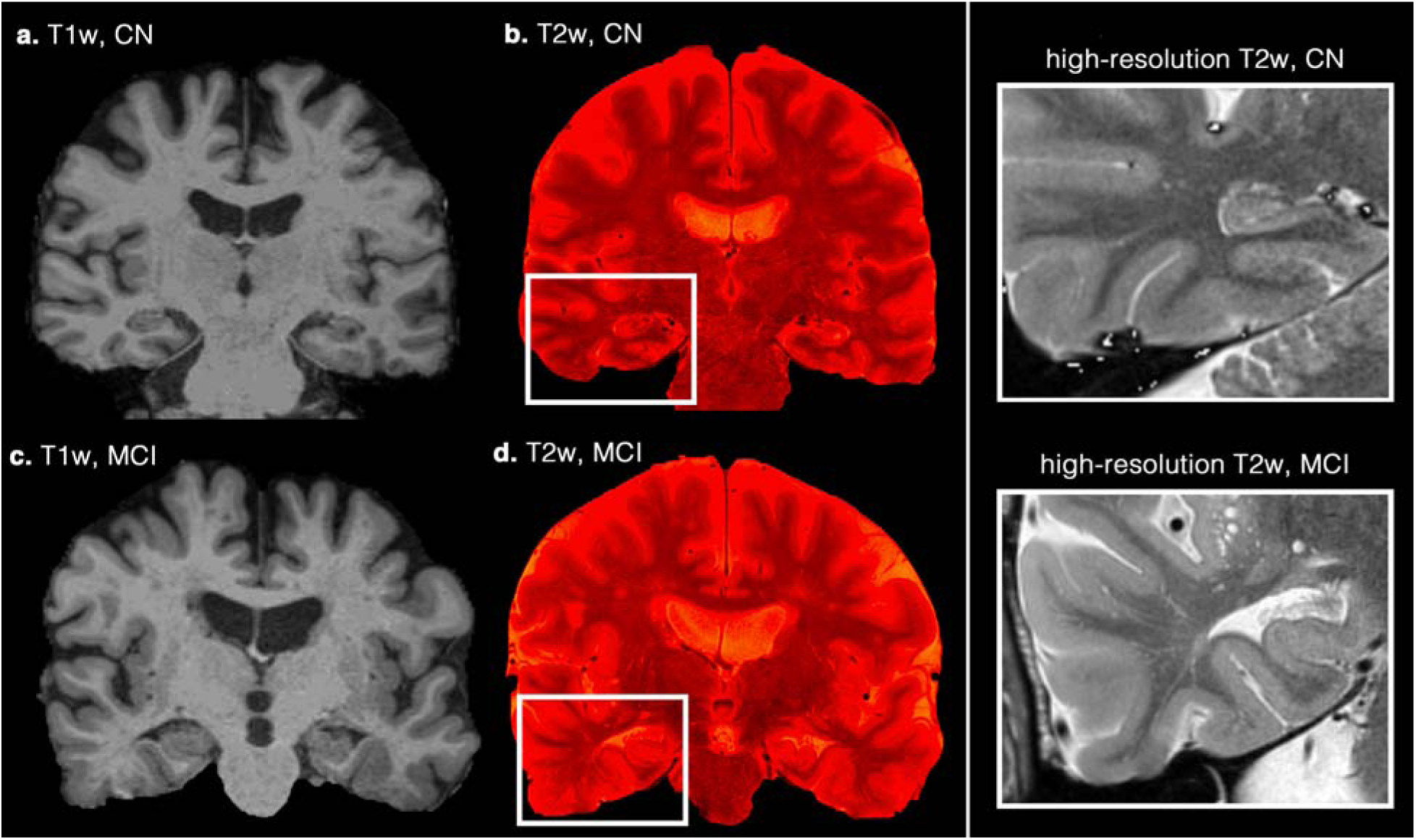
Examples of differing perivascular space (PVS) concentration on a cognitively normal (CN) subject and a patient with mild cognitive impairment (MCI) diagnosis. Coronal T1-weighted images (a and c) are illustrated. High-resolution T2-weighted images were superimposed on them to highlight the PVS (**b** and **d**). Right column figures zoomed into the temporal lobe area, in which increased mean diffusivity and decreased fractional anisotropy are commonly reported.

For additional insight, we further investigated one of the regions with significantly different MD value between CN and MCI from the DTI study, namely the superior fronto-occipital fasciculus. Voxel MD (from DTI), tissue MD (from TTI), and PVS signal fraction of this region are plotted in and compared across CN and MCI in **Figure 7.** Voxel MD was significantly different between CN and MCI when PVS bias was not considered (*p* < 0.001), however, no significant difference was observed in tissue MD. Interestingly, PVS signal fraction appears to be the main feature separating CN and MCI in this region, that is, when it was included as a dependent variable in the regression (*b* = 0.087, *t*(56) = 3.96, *p* < 0.001). When considering all of the regions that we investigated, MD values of DTI were in average 5% higher in the MCI group, but when TTI was used, the MD of the MCI group was in average 1% lower.

**Figure 7.**
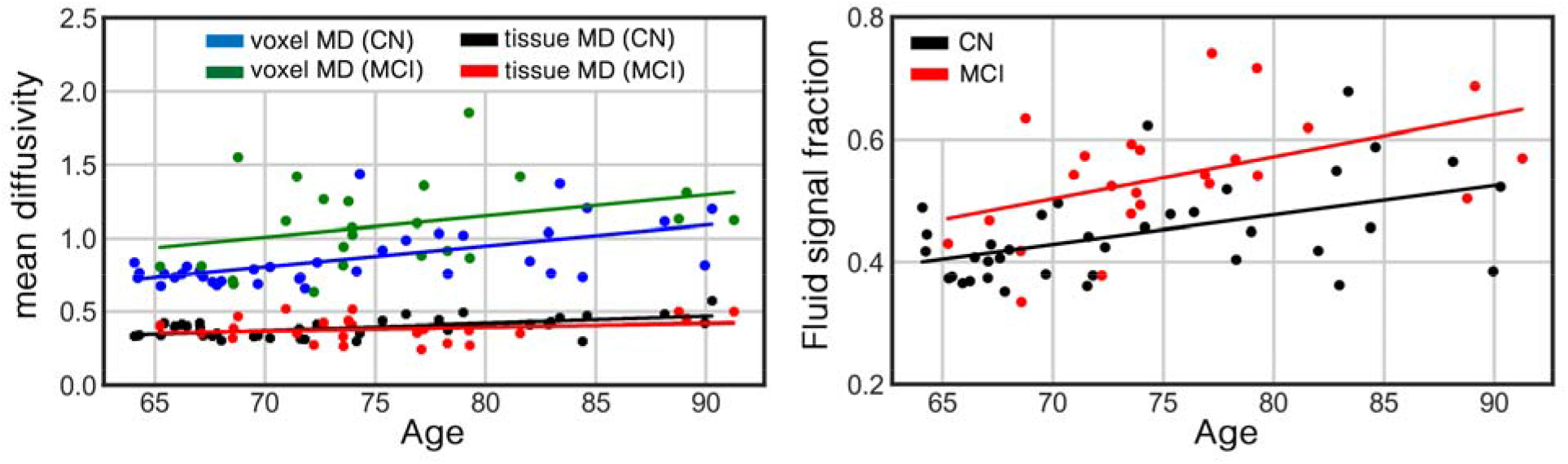
An example of a false discovery of diffusion tensor imaging (DTI) sourced from the PVS bias. An Mean diffusivity (MD) of the left superior fronto-occipital fasciculus of the cognitively normal (CN) group (n=37) and patients with mild cognitive impairment (MCI) diagnosis (n=24) are plotted in left. Comparison are made using both DTI and tissue tensor imaging (TTI) techniques. From DTI eyes, higher MD was significantly associated with the cognitive stage (*p*<0.001). However, when TTI was used, no difference was observed. PVS signal fraction of this region is plotted in right. The PVS signal fraction had significant association with the cognitive stage (*p*<0.001). Age, sex and brain volume were included in the regression analysis as covariates. Diffusivity values are in μm^2^/ms.

Other than PVS fluid, which is physiological in the brain, several pathological features such as cerebral microbleeds and lacunar infarcts, could result in the presence of fluid in the white matter. Such accumulation of fluid, if not modeled, could result to an increased voxel MD. This increase of MD also cannot be interpreted as an increase of white matter tissue MD. Recently, Hoy *et al.* highlighted that “free water compartment” plays an important role in determining the measured values of MD and FA in subjects with MCI (Hoy et al., 2017).

### Limitation

We note that the choices of b-values of the HCP data and the exponential diffusion model are suboptimal for measuring PVS diffusivity, given the non-Gaussian diffusion behavior in high b-values (Assaf et al., 2004; Novikov et al., 2016, 2012; Sepehrband et al., 2017) or due to induced susceptibility of the vasculature network in a monopolar pulse design (Kiselev, 2004; Kiselev and Posse, 1999; Zheng and Price, 2007). Gaussian assumption in high b-value data by itself can bias the diffusivity measures of the diffusion MRI models. Here we used a bi-tensor model (which provides a better fit to the diffusion data than DTI (Pierpaoli and Jones, 2004)) and a commonly used multi-shell design to show that ignoring PVS fluid systematically biases DTI findings. For a robust measurement of PVS diffusion coefficient, we suggest a multi-shell acquisition that includes low b-values (similar to our experimental design) and a more comprehensive model of diffusion profile.

## Acknowledgement

This work was supported by NIH grants: 2P41EB015922-21, 1P01AG052350-01 and USC ADRC 5P50AG005142. The content is solely the responsibility of the authors and does not necessarily represent the official views of the NIH.

## HCP

Data were provided [in part] by the Human Connectome Project, WU-Minn Consortium (Principal Investigators: David Van Essen and Kamil Ugurbil; 1U54MH091657) funded by the 16 NIH Institutes and Centers that support the NIH Blueprint for Neuroscience Research; and by the McDonnell Center for Systems Neuroscience at Washington University.

## ADNI

Data collection and sharing for this project was funded by the Alzheimer’s Disease Neuroimaging Initiative (ADNI) (National Institutes of Health Grant U01 AG024904) and DOD ADNI (Department of Defense award number W81XWH-12-2-0012). ADNI is funded by the National Institute on Aging, the National Institute of Biomedical Imaging and Bioengineering, and through generous contributions from the following: AbbVie, Alzheimer’s Association; Alzheimer’s Drug Discovery Foundation; Araclon Biotech; BioClinica, Inc.; Biogen; Bristol-Myers Squibb Company; CereSpir, Inc.; Cogstate; Eisai Inc.; Elan Pharmaceuticals, Inc.; Eli Lilly and Company; EuroImmun; F. Hoffmann-La Roche Ltd and its affiliated company Genentech, Inc.; Fujirebio; GE Healthcare; IXICO Ltd.; Janssen Alzheimer Immunotherapy Research & Development, LLC.; Johnson & Johnson Pharmaceutical Research & Development LLC.; Lumosity; Lundbeck; Merck & Co., Inc.; Meso Scale Diagnostics, LLC.; NeuroRx Research; Neurotrack Technologies; Novartis Pharmaceuticals Corporation; Pfizer Inc.; Piramal Imaging; Servier; Takeda Pharmaceutical Company; and Transition Therapeutics. The Canadian Institutes of Health Research is providing funds to support ADNI clinical sites in Canada. Private sector contributions are facilitated by the Foundation for the National Institutes of Health (www.fnih.org). The grantee organization is the Northern California Institute for Research and Education, and the study is coordinated by the Alzheimer’s Therapeutic Research Institute at the University of Southern California. ADNI data are disseminated by the Laboratory for Neuro Imaging at the University of Southern California.

## Supplemental Figures

**Supplemental Figure 1.**
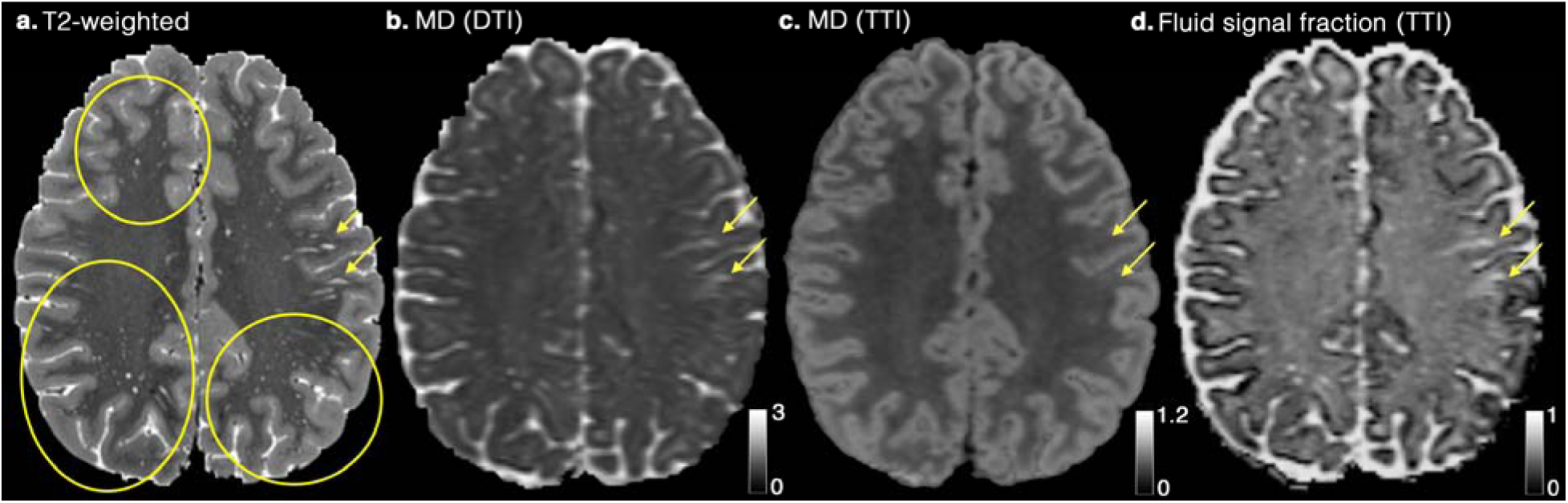
An example of the influence of PVS on mean diffusivity (MD) map. **(a)** T2-weighted image, **(b)** Voxel MD, derived from diffusion tensor imaging (DTI), and **(c)** Tissue MD, derived from tissue tensor imaging (TTI) are shown, **(d)** demonstrates the signal fraction map of the fluid compartment (which includes PVS) derived from diffusion MRI data. Two areas with high PVS concentration are highlighted with yellow arrows. Three regions with high number of visible PVS are also highlighted with yellow ellipsoids. Note that MD from TTI preserves white matter homogeneity and is not affected by PVS contribution, while DTI is significantly affected. MD values are in μm^2^/ms.

**Supplemental Figure 2.**
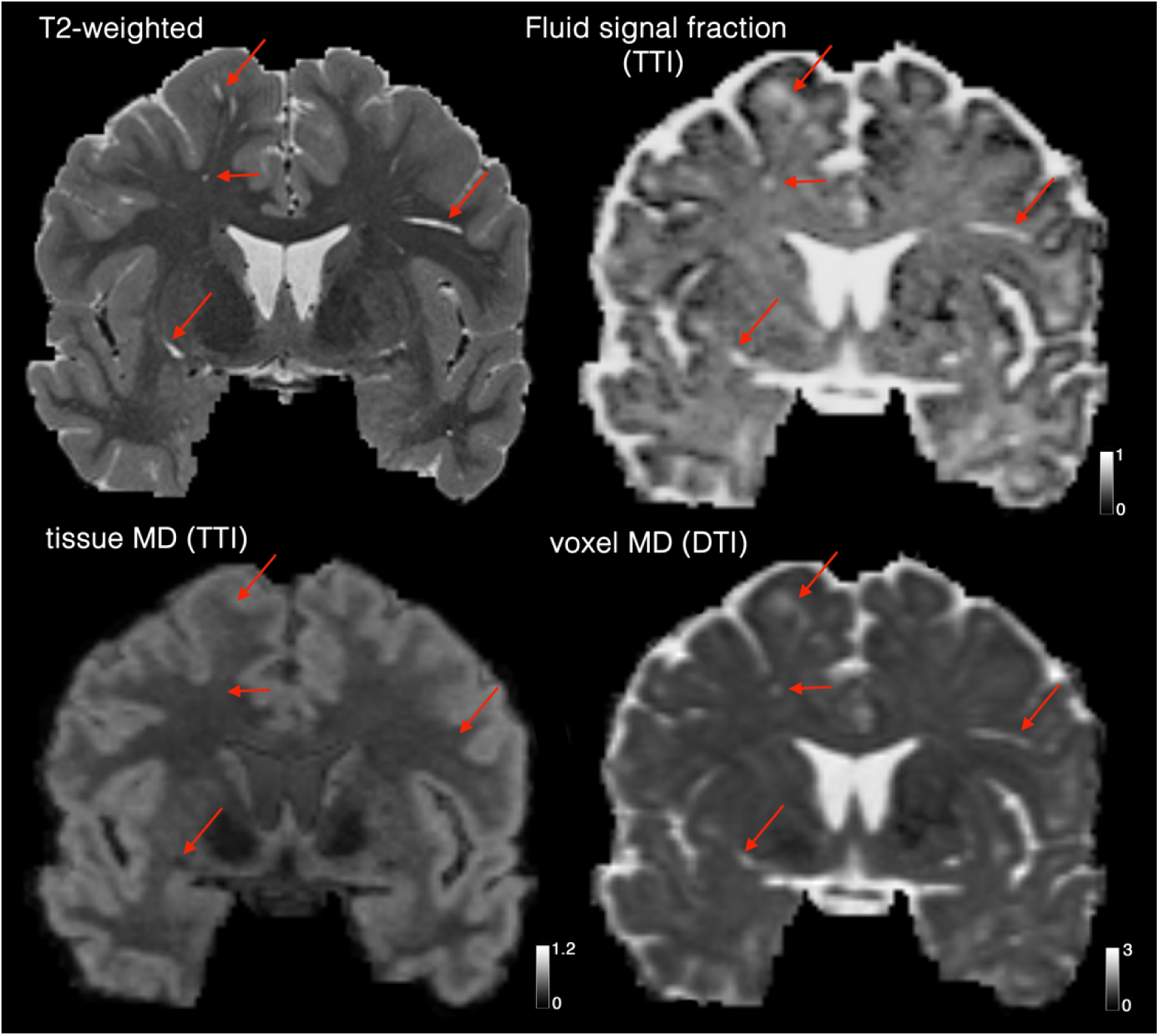
Another example of the influence of PVS on mean diffusivity (MD) map, similar to Supplemental Figure 1.

**Supplemental Figure 3.**
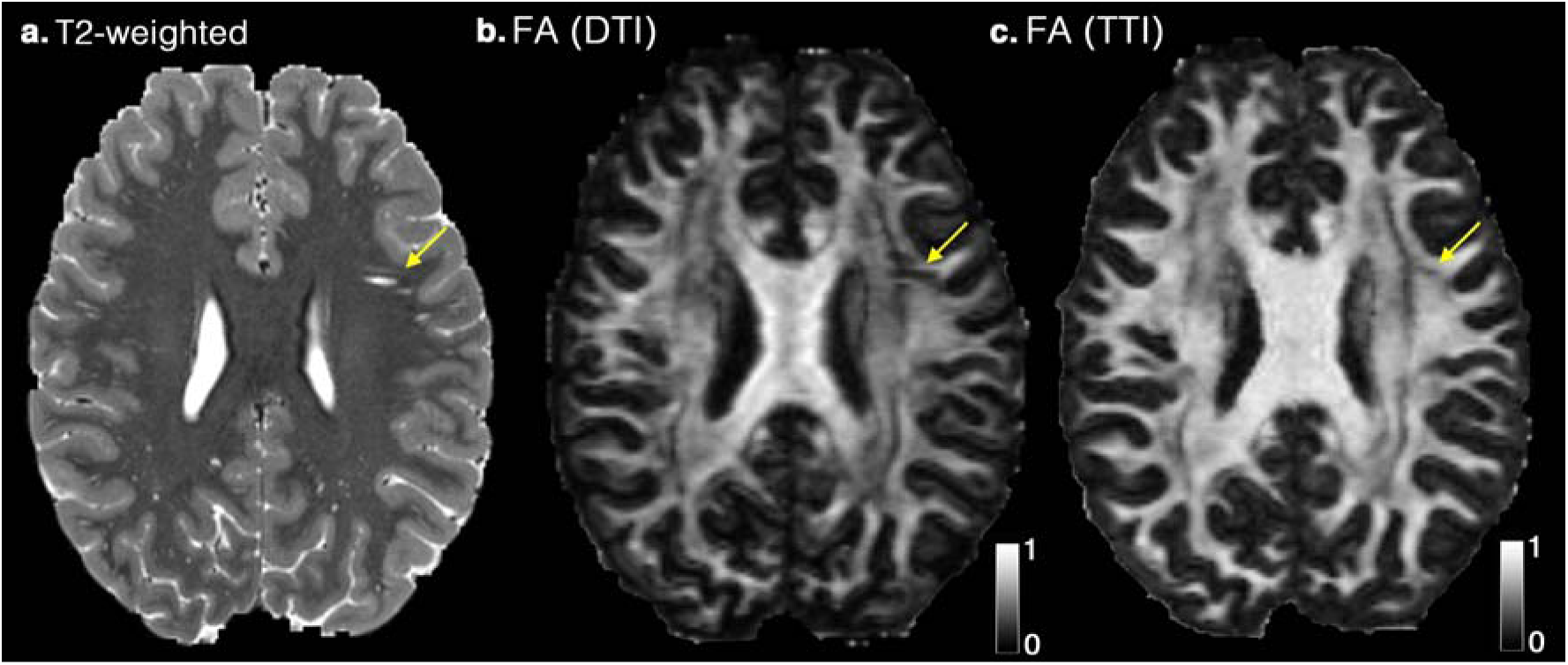
An example of the influence of PVS on fractional anisotropy (FA) map. **(a)** T2-weighted image, **(b)** Voxel FA, derived from diffusion tensor imaging (DTI), and **(c)** Tissue FA, derived from tissue tensor imaging (TTI). An example with high amount of PVS is highlighted with yellow arrows. Note that FA values from TTI are higher throughout the brain.

**Supplemental Figure 4.**
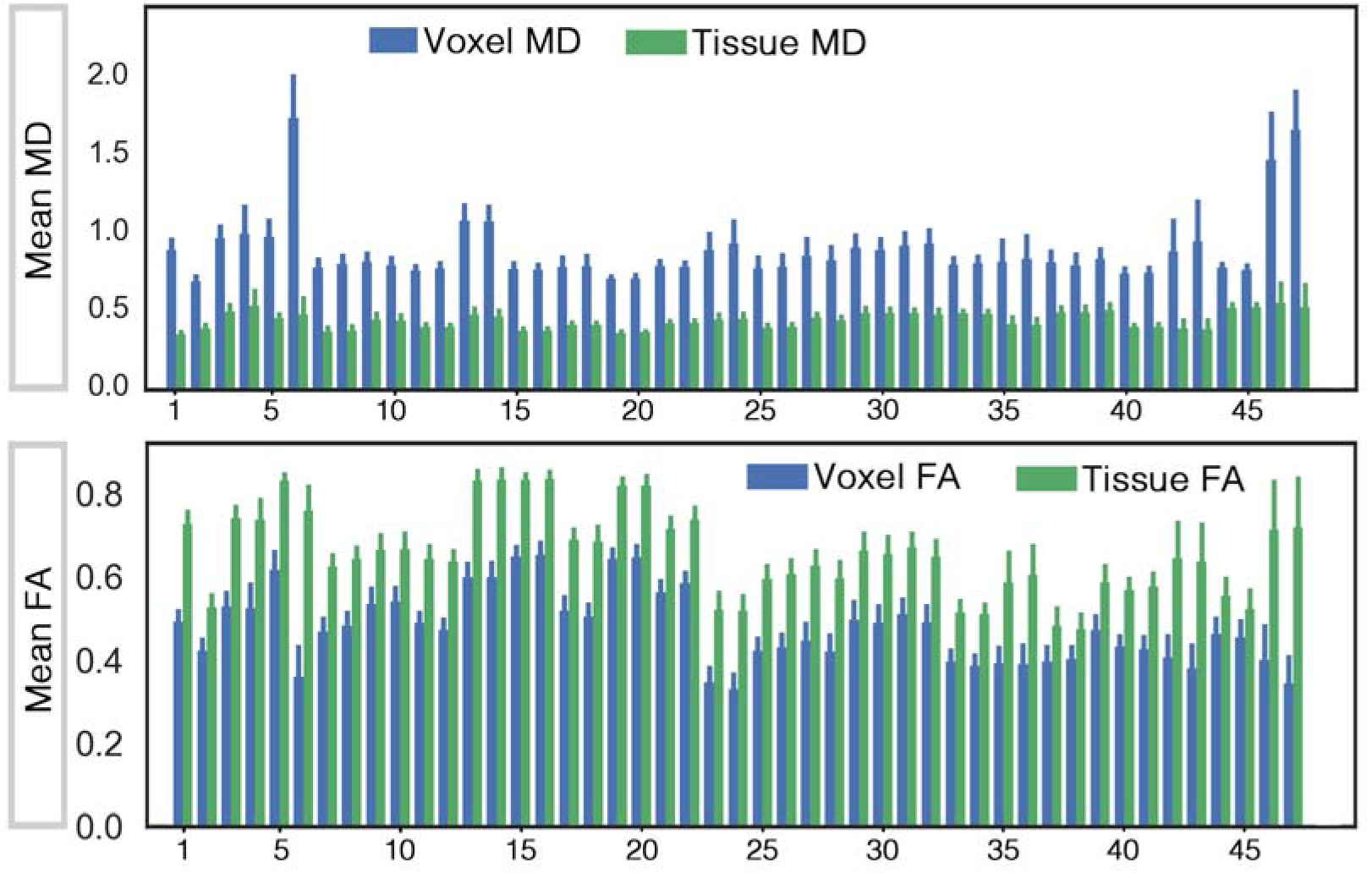
Mean and standard deviation of the mean diffusivity (MD) and fractional anisotropy (FA) across regions of the white matter are plotted (*n*=61, from ADNI-3 dataset). Blue bars are voxel values, derived from diffusion tensor imaging (DTI). Green bars are tissue values, derived from tissue tensor imaging (TTI). Diffusivity values are in μm^2^/ms. Mean values of all regions were significantly different between DTI and TTI, at *p*<0.001. Regions are extracted from John-Hopkins white matter atlas: 1. Middle cerebellar peduncle, 2. Pontine crossing tract (a part of MCP), 3. Genu of corpus callosum, 4. Body of corpus callosum, 5. Splenium of corpus callosum, 6. Fornix (column and body of fornix), 7. Corticospinal tract R, 8. Corticospinal tract L, 9. Medial lemniscus R, 10. Medial lemniscus L, 11. Inferior cerebellar peduncle R, 12. Inferior cerebellar peduncle L, 13. Superior cerebellar peduncle R, 14. Superior cerebellar peduncle L, 15. Cerebral peduncle R, 16. Cerebral peduncle L, 17. Anterior limb of internal capsule R, 18. Anterior limb of internal capsule L, 19. Posterior limb of internal capsule R, 20. Posterior limb of internal capsule L, 21. Retrolenticular part of internal capsule R, 22. Retrolenticular part of internal capsule L, 23. Anterior corona radiata R, 24. Anterior corona radiata L, 25. Superior corona radiata R, 26. Superior corona radiata L, 27. Posterior corona radiata R, 28. Posterior corona radiata L, 29. Posterior thalamic radiation (include optic radiation) R, 30. Posterior thalamic radiation (include optic radiation) L, 31. Sagittal stratum (include inferior longitidinal fasciculus and inferior fronto-occipital fasciculus) R, 32. Sagittal stratum (include inferior longitidinal fasciculus and inferior fronto-occipital fasciculus) L, 33. External capsule R, 34. External capsule L, 35. Cingulum (cingulate gyrus) R, 36. Cingulum (cingulate gyrus) L, 37. Cingulum (hippocampus) R, 38. Cingulum (hippocampus) L, 39. Fornix (cres) Stria terminalis (can not be resolved with current resolution) R, 40. Fornix (cres) Stria terminalis (can not be resolved with current resolution) L, 41. Superior longitudinal fasciculus R, 42. Superior longitudinal fasciculus L, 43. Superior fronto-occipital fasciculus (could be a part of anterior internal capsule) R, 44. Superior fronto-occipital fasciculus (could be a part of anterior internal capsule) L, 45. Uncinate fasciculus R, 46. Uncinate fasciculus L, 47. Tapetum R, 48. Tapetum L.

## Supplemental Notes

**Supplemental Note 1.** Comparing mean FA and mean MD as derived from DTI, DTI-FWE and TTI (this supplemental note accompanies **Figure 5** of the manuscript).

Mean FA comparison across HCP subjects:

- FA values from DTI in PVS area were (*M* = 0.35, *SD* = 0.03)
- FA values from FWE in PVS area were (*M* = 0.46, *SD* = 0.05)
- FA values from TTI in PVS area were (*M* = 0.65, *SD* = 0.04)
- FA from TTI was significantly higher than DTI, *t*(860) = 313.04, *p* < .001, and FWE, *t*(860) = 119.14, *p* < .001, techniques.
- FA from FWE was significantly higher than DTI, *t*(860) = 129.61, *p* < .001
- FA was on average 0.3 higher when measured using TTI compare to DTI (almost two-times higher).
- FA from TTI was significantly correlated with those from DTI *r*(859) = 0.70, *p* < .001
- FA from TTI was significantly correlated with those from FWE *r*(859) = 0.79, *p* < .001
- FA from FWE was significantly correlated with those from DTI *r*(859) = 0.84, *p* < .001

Mean MD comparison across HCP subjects:

- MD values from DTI in PVS area were (*M* = 0.94, *SD* = 0.07)
- MD values from FWE in PVS area were (*M* = 0.39, *SD* = 0.02)
- MD values from TTI in PVS area were (*M* = 0.24, *SD* = 0.02)
- MD from TTI was significantly higher than DTI, *t*(860) = 283.13, *p* < .001, and FWE, *t*(860) = 168.50, *p* < .001, techniques.
- MD from TTI was on average 0.7 higher than that from DTI (almost four time higher).
- MD from FWE was significantly higher than DTI, *t*(860) = 234.86, *p* < .001
- MD from TTI was weakly correlated with those from DTI *r*(859) = −0.06, *p* < .05
- MD from TTI was correlated with those from FWE *r*(859) = 0.25, *p* < .001
- MD from FWE was correlated with those from DTI *r*(859) = 0.11, *p* < .001

**Supplemental Note 2.** Statistically summary from comparing mean diffusivity (MD) values of DTI of cognitively normal (CN) subjects (*n*=37) and patients with mild cognitive impairment (MCI) diagnosis (*n*=24). Eleven regions were significantly different after correcting for multiple comparison using Benjamini–Hochberg procedure with the false discovery rate of 0.1 (age, sex and brain size were included in the regression). Note that none of these differences were significant after incorporating the contribution of the perivascular space.

| White matter region<br>(from John Hopkins WM atlas) | hemisphere | *p*-value | contrast | Is white matter<br>MD different after<br>modeling PVS?<br>(using TTI) |
| --- | --- | --- | --- | --- |
| *Genu of the corpus callosum* | - | 0.023 | MCI > CN | No |
| *Body of the corpus callosum* | - | 0.026 | MCI > CN | No |
| *Splenium of the corpus callosum* | - | 0.011 | MCI > CN | No |
| *Anterior limb of internal capsule* | Left | 0.020 | MCI > CN | No |
| *Anterior corona radiata* | Right | 0.009 | MCI > CN | No |
| *Anterior corona radiata* | Left | 0.004 | MCI > CN | No |
| *Superior corona radiata* | Right | 0.042 | MCI > CN | No |
| *Superior corona radiata* | Left | 0.002 | MCI > CN | No |
| *Posterior corona radiata* | Left | 0.027 | MCI > CN | No |
| *Posterior thalamic radiata* | Right | 0.038 | MCI > CN | No |
| *Posterior thalamic radiata* | Left | 0.036 | MCI > CN | No |
| *Sagittal stratum* | Right | 0.006 | MCI > CN | No |
| *Sagittal stratum* | Left | <0.001 | MCI > CN | No |
| *Cingulum (hippocampus)* | Right | 0.040 | MCI > CN | No |
| *Cingulum (hippocampus)* | Left | 0.019 | MCI > CN | No |
| *Fornix (stria terminalis)* | Right | 0.015 | MCI > CN | No |
| *Superior longitudinal fasciculus* | Right | 0.014 | MCI > CN | No |
| *Superior longitudinal fasciculus* | Left | 0.040 | MCI > CN | No |
| *Superior fronto-occipital fasciculus* | Right | 0.030 | MCI > CN | No |
| *Superior fronto-occipital fasciculus* | Left | <0.001 | MCI > CN | No |
| *Tapetum* | Right | 0.006 | MCI > CN | No |
| *Tapetum* | Left | 0.003 | MCI > CN | No |

